# The ExtRA Capacity Test: Reliability and validity study of a new tool for assessing shoulder muscle performance

**DOI:** 10.1101/2022.06.23.496982

**Authors:** Harry Ford, Jeremy Lewis, Vasileios Tyros, Marco Davare, Daniel Low, Aliah Shaheen

**Affiliations:** Brunel University London, College of Health, Medicine and Life Sciences; Central London Community Healthcare National Health Service Trust; PCRF Pirbright; King’s College London, Life Sciences and Medicine; Brunel University London, Sports, Health and Exercise Sciences

## Abstract

**Objectives:** The primary objective was to evaluate the reliability of a new tool for assessing shoulder muscle performance: The ExtRA Capacity Test. The secondary objective was to assess whether this tool was a valid measure for assessing shoulder strength.

**Methods:** The ExtRA Capacity test involves two measures: maximal scapular plane lateral raises to 90° abduction with 2.5kg of external load and the maximal number of prone lying unsupported external rotations with the shoulder at 90° abduction. Both capacity tests are completed to a metronome set to 30 beats per minute 20 asymptomatic participants were sampled on 2 separate sessions, 1 week apart. The ExtRA Capacity Test was completed by the lead researcher and an independent physiotherapist. Shoulder strength was also measured using isokinetic dynamometry.

**Results:** The test showed excellent inter-rater reliability (mean abduction ICC= 0.969, mean external rotation ICC= 0.822, with a 95% CI). Mean intra-rater variability was 3.96± 4.09 for the abduction measure and 1.70± 1.17 for the external rotation measure. Validity was calculated using Pearson correlation coefficient. The abduction measure showed good/ moderate correlation for the majority of strength measurements taken using isokinetic dynamometry however the external rotation capacity test did not correlate closely to isokinetic dynamometry strength measures.

**Conclusion:** The abduction component of the ExtRA Capacity Test is a suitable measure for assessing shoulder strength in clinical practice. The external rotation measure is of suitable reliability however if used in clinical practice, it should not be used to assess shoulder strength, instead it may be suitable to assess movement control of the shoulder.

**Summary:** Various methods of measuring shoulder strength exist, ranging from cost free, relatively inaccurate methods to costly, complex methods which are of high reliability and validity but are challenging to use in a fast-pace, clinical environment. Objective outcome measures are used within a rehabilitation setting however at present there is no upper limb muscle performance test that is suitable to use on all people, regardless of strength or fitness level. Capacity testing of movements provides a functional, insight into strength specific to a real world/ sporting environment with lower limb capacity tests providing clear objective baselines that can be used for goal setting and providing return to play criteria following injury. This study proves the reliability and validity of the ExtRA Capacity Test which is a measure of shoulder muscle performance, suitable for people of all physical activity levels and upper limb strength.

## 1. Introduction

Reduction in muscle strength can occur as a result of injury, trauma, kinesephobia or as a result of disuse due to lifestyle or following surgery (Campbell et al., 2013; Greenhaff. 2006). There is conflicting evidence surrounding whether shoulder strength should be used as a predictor of atraumatic shoulder injury (Asker et al., 2018; Stuelcken et al., 2008; Bagordo et al., 2020). Despite this, shoulder strength measures are often used in conjunction with range of movement and functional outcome measures to judge when a return to sport following shoulder injury is appropriate (Cools et al., 2020). There are various methods of measuring shoulder strength. Isokinetic Dynamometry (IKD) has become an increasingly popular assessment tool used in exercise science and sports medicine and is widely recognised as the gold standard for measuring muscle strength and muscle endurance through a specified range of movement (Osternig, 1986; Bagordo et al., 2020). The high cost of IKD means that is it rarely used in clinical practice with other more cost-effective methods being favoured including hand held dynamometry and manual muscle testing. While hand held dynamometry has been found to provide good to excellent intra- and inter-tester reliability for the measurement of shoulder strength, it only is able to provide values for isometric muscle strength which does not correspond highly to the functional demands of the shoulder (Dollings et al., 2012; Cools et al., 2020). On the other hand, while manual muscle testing is able to measure shoulder strength through range, it lacks precision with side-to-side differences in strength only being detectable when muscle strength was less than 75-85% of that on the contralateral side (Nagatomi et al., 2017).

There are many mechanisms which play a role in the development and recurrence of shoulder pain (Lewis, 2016; Littlewood et al.,2019). The biomedical model is the most widely researched of these with significant evidence suggesting that exercise-based interventions can be good for treating shoulder pain however at present there is no existing upper limb muscle performance test that is suitable to use on all people, regardless of strength or fitness level (Holmgren et al., 2012; Lewis, 2016; Littlewood et al., 2019; Richardson et al., 2020). Capacity testing of movements may provide a more functional, accurate insight into shoulder strength as it replicates movements that may be completed in a real world/ sporting environment and also gives a clear numerical value that can provide a clear objective outcome measure without the need of costly, laboratory equipment (Kollock et al., 2015). A body of research using a maximum repetition method to monitor strength already exists for the lower limb, with the emergence of normative values providing clear objective baselines which can be used in for goal setting and providing return to play criteria following injury (Culvenor et al., 2016; Hébert-Losier et al., 2017).

## 2. Methods

### 2.1 Participants

A convenience sample of 20 participants aged between 19-58 (13 men and 7 women) participated in the study. The participants were recruited through a variety of different methods: social media, posters around university campus, the universities webpage and the opportunity to participate in this study was also discussed in undergraduate lectures. Participants were excluded from the assessment if they had any conditions that might affect the muscle strength: systemic illness, pregnancy, cervical/ shoulder pain during rest or during active movement, history of cervical/ shoulder pain or treatment to this region over the past 12 months, history of spinal or upper limb surgery or history of spinal or upper limb fractures. The study protocol was approved by the Brunel University College of Health, Medicine and Life Sciences Research Ethics Committee with ethical approval number 31585-LR-Oct/2021-34399-2. All participants were informed about the study procedures and gave written informed consent prior to participating in the study.

### 2.2 The ExtRA Capacity Test

The ExtRA (External Rotation/ Abduction) Capacity Test involves measuring the maximum number of repetitions performed before fatigue or a loss of form of scapular plane shoulder abduction with 2.5kg of external load and prone lying external rotation.

For the abduction capacity test, participants stood with their feet shoulder width apart with their back to a wall and were asked to abduct their arm to 90° in the scapular plane while holding a 2.5kg dumbbell. The rate of movement was controlled by a ‘30 beats per minute metronome’ with one beat signalling the concentric phase of the movement and the other beat signalling the eccentric phase of the movement. The test was terminated when subjects could: (1) could no longer perform a full shoulder abduction to 90° with the external load applied; (2) could no longer maintain the set pace; or (3) could no longer control the movement or trunk alignment.

Prone lying external rotation capacity measurements were taken with the participant in prone lying on a flat treatment couch with their arm out at 90° abduction, elbow at 90° flexion and their forearm pronated with the palm facing the floor. The examiner initially supported their upper arm and passively moved their shoulder into maximal external rotation, marking the point that the head of the ulnar reaches in this position. The participant was then instructed to perform this full movement actively, with their upper arm unsupported. During the assessment, the examiner was instructed to use a single finger on each hand, one hand marking the olecranon and the other to mark the position of the head of ulnar when the shoulder is in maximal external rotation. Similar to the shoulder abduction capacity test, the pace of the movement was controlled by a ‘30 per minute metronome’. The test was terminated when subjects could: (1) could no longer actively perform the movement to their full passive range (ulnar head failed to reach maximal passive position marked by the examiner); (2) could no longer maintain the set pace; or (3) could no longer control of the movement (olecranon moves more than 1cm away from the position marked by the examiner in any direction, demonstrated in Figure 3).

**Figure 1:**
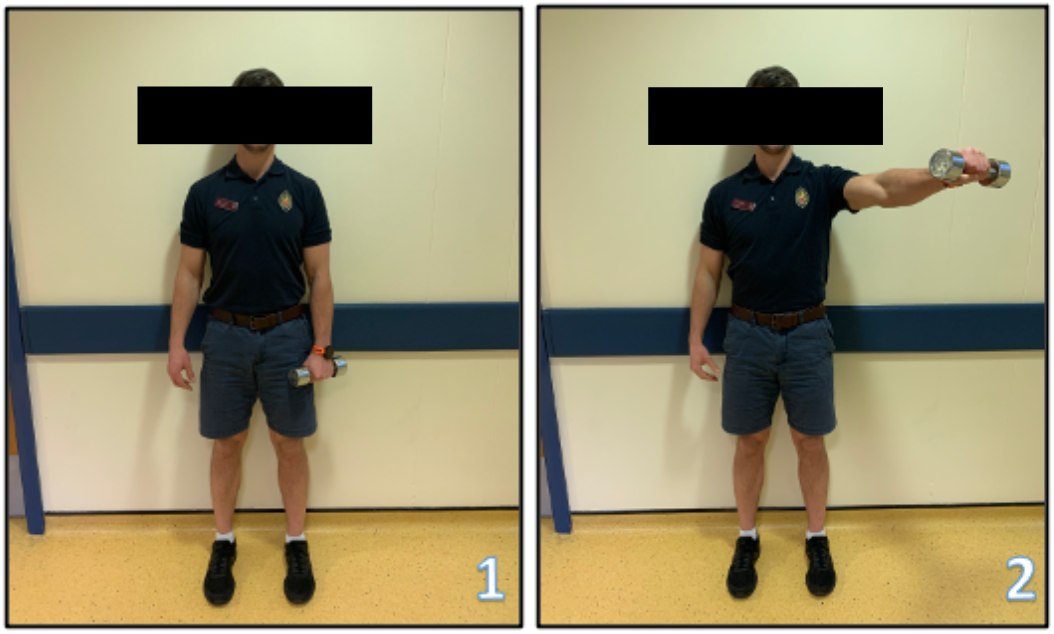
Model demonstrating the ExtRA Abduction Capacity Test. Picture 1: Starting position of capacity test. Picture 2: Participant raised 2.5kg kettlebell to 90degrees abduction in scapular plane.

**Figure 2:**
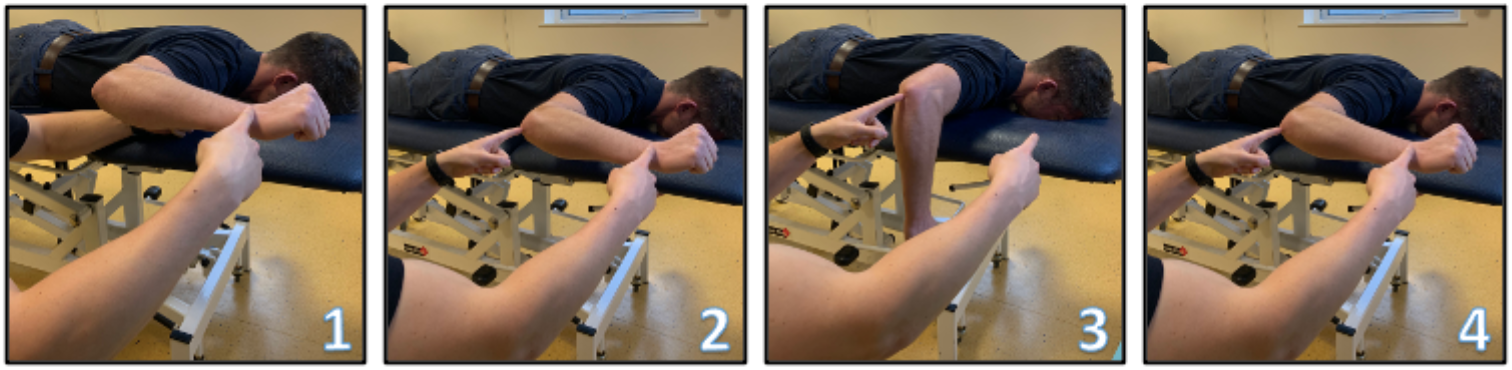
Model demonstrating the ExtRA External Rotation Capacity Test. Picture 1: Examiner supporting participants upper arm so it is in line with their upper back and noting position of the head of the ulnar when in maximal external rotation. Picture 2: Examiner marking olecranon Picture 3: Participant asked to internally rotate arm around point marked on olecranon. Picture 3: Participant rotating arm back up until head of ulnar makes contact with examiners hand.

**Figure 3:**
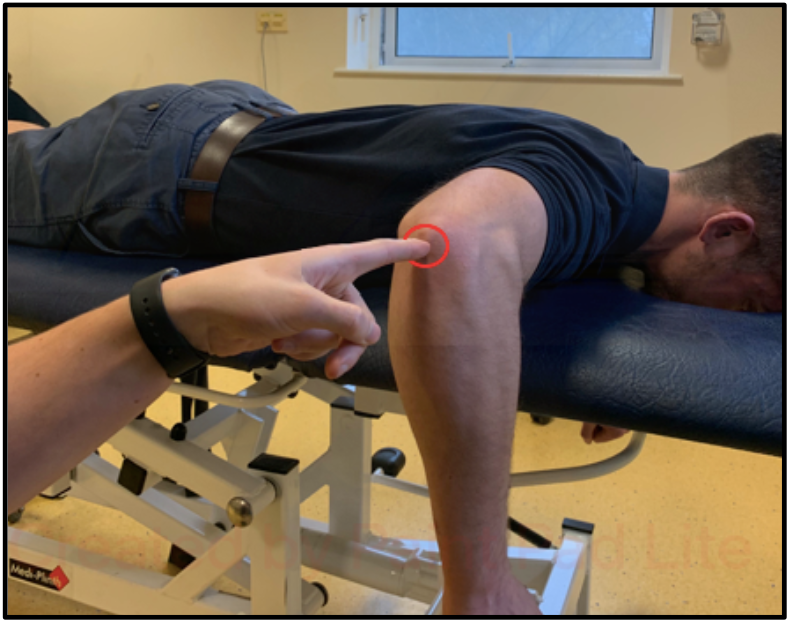
Demonstration of the area that the examiners index finger, marking the olecranon should stay in during the ExtRA External Rotation Capacity Test. This is visualised.

### 2.3 Procedures

All the participants completed two testing sessions on separate days, 1 week apart. The first session included the ExtRA Capacity Test being taken by the lead investigator as well as shoulder strength being assessed through IKD. Participants were given 30 minutes rest between the two strength tests where they were instructed to wait in a waiting room and have a drink of water. Half the participants completed the ExtRA Capacity test before IKD and the other half completed the IKD testing before the ExtRA Capacity test.

Participants were taken through a structured warmup by the lead researcher prior to undergoing the strength testing. The warm up was comprised of nine exercises: range of movement work for the shoulder and cervical region, followed by resistance banded shoulder abductions, resistance banded shoulder external rotation and resistance banded overhead press (Wattanaprakornkul et al, 2011a; Wattanaprakornkul et al., 2011b).

Isokinetic testing of both shoulders was performed using a Biodex System 4 Isokinetic System (Harding et al., 2015). Isokinetic testing was completed for shoulder abduction capacity in the scapular plane (from 0deg-90deg abduction) as well as external rotation capacity at 90degrees abduction (from maximal external rotation-0degrees internal rotation). In order to assess external rotation and abduction capacity, the Biodex System 4 was set using ‘Passive Mode’ at 30°/s for 20 repetitions as this is the same rate as the ExtRA Capacity Test (Harding et al., 2015). For the external rotation capacity testing, participants were instructed to maximally push into external rotation and then to maximally resist the lever returning to the starting position. This allowed for both concentric and eccentric external rotation strength values to be calculated. A similar instruction was given when monitoring abduction capacity: pull up as hard as possible and try to prevent the arm moving down. Prior to maximal testing of each movement, the participant was allowed to complete 5 practice repetitions using the Biodex System 4 Isokinetic System, 2 minutes was then be left prior to the maximal test. There was a 5-minute rest between isokinetic testing in different positions. Monitoring the output of the 20 repetitions allowed for 10 different values to be calculated for both shoulder abduction and shoulder external rotation: Concentric/ eccentric values of peak torque, average peak torque, total work, work fatigue and average power (Tonin et al., 2013; Ellenbecker and Roetert 1999; Bagordo et al., 2020).

During the second testing session, the ExtRA Capacity Test was repeated by the lead researcher as well as an independent physiotherapist who was following the same instructions. Similar to the first testing session, there was a 30-minute rest between capacity tests and half of the participants were assessed by the lead investigator before the independent physiotherapist and vice versa. Scores recorded by the lead investigator were compared to scores taken by the independent physiotherapist in order to assess interrater reliability. The values recorded by the lead investigator were also compared to those that they gathered 1 week earlier so intrarater reliability can be assessed.

### 2.4 Statistical analysis

A sample size of 20 was selected as it is within the range of other, recent, reliability tests looking at objective outcome measures: 13, 16 and 40 respectively (Dos’ Santos et al., 2018; Dingenen et al., 2019; Thongchoomsin et al., 2020). Pearson correlation coefficients were used to examine the relationship between shoulder strength values obtained by IKD and the ExtRA Capacity Test taken by the lead researcher (Koo and Li, 2016).

ICC was used in conjunction with standard deviation measures to order to assess the ExtRA measurement reliability, interobserver variability as well as ‘between session variability’. Significance level was set at p < 0.05 for all analyses (Obilior and Amadi, 2018).

## 3. Results

Of the 20 participants who were sampled, all 20 of them reported being right-handed. There were 13 males and 7 females with a mean age of 28 (SD= 13.28). Participant weight ranged from 51.5kg-112.1kg with a mean of 78.91kg ± 18.14. Mean height of participants was 173.92cm (SD= 11.58) with a range of 157.5cm-196cm. Nineteen of the 20 participants completed all of the sampling. One participant however sustained a shoulder injury between the 2 sampling sessions so only their dominant arms strength values were gathered in both sessions.

### 3.1 Inter Observer and between session reliability/ variability

The ICC values calculated demonstrate excellent inter rater reliability for the abduction capacity test for both the dominant and non-dominant arm (0.980 and 0.958 respectively). Good inter rater reliability was found for both the dominant and non-dominant external rotation measures with an ICC of 0.874 for the dominant arm and 0.769 for the non-dominant arm (Koo and Li, 2016).

Excellent intra rater reliability was reported for dominant and non-dominant abduction measure as well as for the non-dominant arm external rotation measure (ICC=0.966, 0.959,0.936). Good reliability was reported for the dominant arm external rotation reliability (ICC= 0.890) (Koo and Li, 2016).

The between session variability was higher for the abduction capacity test when compared to the external rotation capacity test (4.38 ± 4.09/ 2.05 ± 1.2 for the dominant arm and 3.61 ± 4.09/ 1.34 ± 1.13 for the non-dominant arm). It is worth noting that the mean scores were considerably higher than for the abduction measure than the external rotation capacity test.

### 3.3 ExtRA Capacity Tests validity for measuring shoulder strength

The Pearson correlation coefficient calculations are documented in Table 2. When assessing the correlation between the ExtRA abduction capacity test and strength measures gathered via IKD, only 7 out of 40 measurements were found to be of insufficient significance for the findings to be meaningful (>0.05) (Obilior and Amadi, 2018). Of those 33 measurements of sufficient significance, 6 strength values calculated via IKD were considered to have a strong correlation to the ExtRA abduction capacity test (including both concentric abduction total work and average concentric abduction power for the dominant and non-dominant arm). 26 of the 40 IKD strength measures demonstrated moderate correlation to the abduction strength measure and only one showed low correlation (Munro, 2005).

**Table 1:** Table demonstrating inter and intra rater mean variability and ICC for each of the ExtRA measures.

|  | Inter Rater Reliability/ Variability |  |  |  | Between Session Reliability/ Variability |  |  |  |
| --- | --- | --- | --- | --- | --- | --- | --- | --- |
| ExtRA Measure | Lead Researcher Mean | Independent Physiotherapist Mean | Mean Inter Observer Variability | ICC (95% CI Lower Bound, Upper Bound) | Session 1 Mean | Session 2 Mean | Mean Between Session Variability | ICC (95% CI Lower Bound, Upper Bound) |
| Dominant Abduction | 47 ± 23.8 | 45.3 ± 22.88 | 3.82 ± 2.75 | 0.980 (0.949, 0.992) | 47.05 ± 24.08 | 53.1 ± 23.9 | 4.38 ± 4.09 | 0.966 (0.841, 0.973) |
| Non-Dominant Abduction | 41.9 ± 20.67 | 40.95 ± 20.21 | 6.33 ± 4.91 | 0.958 (0.894, 0.983) | 40.11 ± 21.68 | 44.67 ± 21.12 | 3.61 ± 4.09 | 0.959 (0.809, 0.969) |
| Dominant External Rotation | 15.55 ± 6.53 | 15.7 ± 7.6 | 2.44 ± 2.24 | 0.874 (0.681, 0.95) | 15.05 ± 6.1 | 15.4 ± 5.89 | 2.05 ± 1.2 | 0.890 (0.564, 0.916) |
| Non-Dominant External Rotation | 14.65 ± 4.96 | 13.35 ± 5.28 | 2.55 ± 1.93 | 0.769 (0.416, 0.909) | 13.37 ± 5.0 | 14.67 ± 5.02 | 1.34 ± 1.13 | 0.936 (0.715, 0.952) |

**Table 2:** Table demonstrating the correlation between the ExtRA Capacity Test and strength measured obtained by Isokinetic Dynamometry

|  |  | Concentric<br>Peak<br>Torque | Concentric<br>Average Peak<br>Torque | Concentric<br>Total<br>Work | Concentric<br>Work<br>Fatigue | Concentric<br>Average<br>Power | Eccentric<br>Peak<br>Torque | Eccentric<br>Average Peak<br>Torque | Eccentric<br>Total<br>Work | Eccentric<br>Work<br>Fatigue | Eccentric<br>Average<br>Power |
| --- | --- | --- | --- | --- | --- | --- | --- | --- | --- | --- | --- |
| Abduction Capacity Test |  |  |  |  |  |  |  |  |  |  |  |
|  |  | Isokinetic Dynamometry Measure: Abduction |  |  |  |  |  |  |  |  |  |
| Dominant | Pearson Correlation | 0.616 | 0.607 | 0.817 | 0.268 | 0.752 | 0.556 | 0.533 | 0.576 | 0.135 | 0.514 |
|  | Sig. (2-tailed) | 0.004 | 0.005 | 0 | 0.254 | 0 | 0.011 | 0.016 | 0.008 | 0.57 | 0.02 |
| Non-Dominant | Pearson Correlation | 0.616 | 0.718 | 0.797 | 0.338 | 0.739 | 0.584 | 0.542 | 0.601 | 0.351 | 0.533 |
|  | Sig. (2-tailed) | 0.005 | 0.001 | 0 | 0.157 | 0 | 0.009 | 0.017 | 0.007 | 0.14 | 0.019 |
|  |  | Isokinetic Dynamometry Measure: External Rotation |  |  |  |  |  |  |  |  |  |
| Dominant | Pearson Correlation | 0.764 | 0.69 | 0.661 | 0.413 | 0.689 | 0.698 | 0.68 | 0.531 | -0.07 | 0.583 |
|  | Sig. (2-tailed) | 0 | 0.001 | 0.002 | 0.07 | 0.001 | 0.001 | 0.001 | 0.016 | 0.768 | 0.007 |
| Non-Dominant | Pearson Correlation | 0.692 | 0.679 | 0.623 | 0.487 | 0.615 | 0.596 | 0.509 | 0.514 | 0.263 | 0.519 |
|  | Sig. (2-tailed) | 0.001 | 0.001 | 0.004 | 0.034 | 0.005 | 0.007 | 0.026 | 0.024 | 0.277 | 0.023 |

| External Rotation Capacity Test |  |  |  |  |  |  |  |  |  |  |  |
| --- | --- | --- | --- | --- | --- | --- | --- | --- | --- | --- | --- |
|  |  | Isokinetic Dynamometry Measure: Abduction |  |  |  |  |  |  |  |  |  |
| Dominant | Pearson Correlation | 0.108 | 0.074 | 0.234 | 0.342 | 0.211 | -0.026 | -0.038 | 0.186 | 0.206 | 0.107 |
|  | Sig. (2-tailed) | 0.652 | 0.756 | 0.321 | 0.14 | 0.373 | 0.914 | 0.874 | 0.432 | 0.383 | 0.653 |
| Non-Dominant | Pearson Correlation | 0.309 | 0.373 | 0.476 | 0.199 | 0.403 | 0.268 | 0.245 | 0.334 | 0.253 | 0.236 |
|  | Sig. (2-tailed) | 0.197 | 0.116 | 0.04 | 0.415 | 0.087 | 0.267 | 0.312 | 0.163 | 0.295 | 0.331 |
|  |  | Isokinetic Dynamometry Measure: External Rotation |  |  |  |  |  |  |  |  |  |
| Dominant | Pearson Correlation | 0.346 | 0.29 | 0.204 | 0.35 | 0.171 | 0.207 | 0.159 | 0.193 | 0.285 | 0.159 |
|  | Sig. (2-tailed) | 0.135 | 0.214 | 0.388 | 0.13 | 0.47 | 0.382 | 0.504 | 0.415 | 0.223 | 0.502 |
| Non-Dominant | Pearson Correlation | 0.482 | 0.393 | 0.328 | 0.324 | 0.316 | 0.322 | 0.26 | 0.268 | 0.153 | 0.259 |
|  | Sig. (2-tailed) | 0.037 | 0.096 | 0.17 | 0.177 | 0.188 | 0.179 | 0.282 | 0.267 | 0.531 | 0.285 |

| Significance < 0.05 | Very Low Correlation (0-0.25) | Low correlation (0.26-0.49) | Moderate correlation (0.5-0.69) | Strong correlation (0.7-0.89) | Very strong correlation (0.9-1.0) |
| --- | --- | --- | --- | --- | --- |

**Table 3:** Between session variability for participants who completed IKD before/ after ExtRA capacity test.

| ExtRA Measure | Mean Between Session Variability |  | ICC (95% CI Lower Bound, Upper Bound) |  |
| --- | --- | --- | --- | --- |
|  | IKD Completed First | IKD Completed Second | IKD Completed First | IKD Completed Second |
| Dominant Abduction | 6.08 ± 4.95 | 2.69 ± 2.13 | 0.923 (0.692, 0.981) | 0.99 (0.919, 0.995) |
| Non- Dominant Abduction | 4.81 ± 5.82 | 2.28 ± 1.36 | 0.92 (0.677, 0.98) | 0.991 (0.925, 0.996) |
| Dominant External Rotation | 2.26 ± 1.28 | 1.84 ± 1.16 | 0.767 (0.63,0.942) | 0.929 (0.556, 0.965) |
| Non-Dominant External Rotation | 1.77 ± 1.38 | 0.86 ± 0.47 | 0.887 (0.546, 0.972) | 0.981 (0.843, 0.991) |

The ExtRA external rotation capacity test correlated much less closely to shoulder strength values obtained using IKD with 37 of the 40 measurements proving to be of insufficient significance for the findings to be meaningful. Of the 3 strength measures that were found to have a significance of <0.05, all 3 measures demonstrated a low correlation.

## 4. Discussion

This current study was carried out to determine the reliability and validity of the ExtRA capacity test. The ExtRA capacity test involves assessing maximal repetitions of loaded abduction and external rotation capacity as these are movements that reflect shoulder use in real world/ sporting environments (Chu et al., 2016; Wattanaprakornkul et al, 2011a; Wattanaprakornkul et al., 2011b). It is also a test that can easily be performed in a clinical setting without requirement for complex, costly equipment.

The study findings supported that the ExtRA capacity test has satisfactory inter observer reliability. For both the abduction and external rotation capacity test the reliability was greater on testing of the dominant arm as well as a reduced mean interobserver variability. A possible reason for this observation may be due to the impact that short term learning effect has on the outcome of the capacity test. Short term learning effect of the ExtRA Capacity Test may have had a smaller effect on the dominant arm when compared to the non-dominant arm due to increased familiarisation that the individual has performing skill-based tasks with the dominant limb (Gonzalez et al., 2015; Schweiger et al., 2021).

The study findings also demonstrate that the ExtRA capacity test has a good intra-rater reliability and relatively small between-session variability with three out of four of the capacity tests having an excellent reliability and the other having a good reliability (Koo and Li, 2016). This between session reliability is comparable to other measures of shoulder strength/ function including hand held dynamometry and the Closed Kinetic Chain Upper Extremity Stability test (CKCUES test) (Awatani et al., 2016; Tucci et al., 2014; de Oliveira et al., 2017).

This study’s findings suggest that the abduction component of the Extra Capacity Test correlates closely to the strength values obtained through using IKD; both for concentric and eccentric abduction and external rotation strength. These findings suggest that the ExtRA abduction capacity test is of superior accuracy to existing cost free, clinic-based measures of shoulder strength (Nagatomi et al., 2017).

Contrastingly, the external rotation component of the ExtRA capacity test was found to be very poorly correlated to shoulder strength values gathered using IKD. This is likely due to the fact that this test was not purely a measure of shoulder strength as the test would be terminated if the participant either lost control of the movement or was to reach fatigue. This meant that a lack of movement control could lead to a participant who has high external rotation strength output when the upper arm was supported to receive a poor ExtRA rotation score.

## 5. Clinical Implications

Various methods of measuring shoulder strength exist (Awatani et al., 2016; Bagordo et al., 2020; Nagatomi et al., 2017). Within a physiotherapy setting shoulder strength measurements are used to assess shoulder function, monitor response to an intervention as well as guide a return to play following an injury (Chaconas et al., 2017; Hurley et al., 2021).

This reliability and validity study suggests that the ExtRA Capacity Test may be of high clinical use in a rehabilitation setting with the abduction measure providing a simple, reliable method for assessing shoulder strength. While found to be of suitable reliability, the ExtRA external rotation measure is not appropriate to assess shoulder strength. This measure may still be of clinical use as it has potential to assess muscular control of the shoulder (Worsley et al., 2013). This is relevant as muscular control training has been found to reduce pain and increase function for patients with shoulder pain (Bae et al., 2011; Worsley et al., 2013; Roy et al., 2009).

Future research should aim to gather normative values for both the ExtRA abduction and external rotation capacity test for both active and inactive asymptomatic individuals of both sexes and a range of ages. This data will be of use for future research into the impact that shoulder strength has on the long term prognosis of shoulder pain. It will enable comparisons to be drawn between long term shoulder function in previously symptomatic individuals who did/ did not achieve the normative values based on their respective age and sex. Normative values will also be of useful when goal setting and monitoring response to treatment in a clinical setting. Similar data already exists for lower limb capacity tests which provides clear objective baselines which can be used in for goal setting and providing return to play criteria following injury (Culvenor et al., 2016; Hébert-Losier et al., 2017).

## 6. Limitations

Limitations to this study should be noted. First, in testing session 1, half of the participants completed the IKD strength measure before the ExtRA capacity test and the other half completed the ExtRA capacity test first. While it was ensured that there was 30 minutes rest between the 2 strength tests in order to eliminate the effect of fatigue, results suggests that this was not long enough as subgroup analysis indicates that between session variability was higher and the ICC was lower in the group that completed the IKD before the ExtRA capacity test in session 1. This suggests that the final intrarater reliability values reported were underestimated by the design of study.

Another limitation to this study was that participant activity in the days before the initial testing session and the week between the two testing sessions was not controlled. Participants were free to participate in all activities of their daily lives which meant that some would have participated in strenuous exercise/ activity in the days prior to testing which may have led them to be fatigued at the point of testing. Activity level not being controlled between testing sessions was also found to be an issue as one participant sustained a shoulder injury during contact sport between the two testing sessions which meant that only strength values for their dominant arm were gathered in both sessions.

